# No evidence that lulworthioid fungi are dark septate endophytes in the roots of the dominant Mediterranean seagrass *Posidonia oceanica*

**DOI:** 10.1101/2021.05.05.442788

**Authors:** Martin Vohník

## Abstract

A previous study from Sicily, Italy indicated that the dominant Mediterranean seagrass *Posidonia oceanica* forms a dark septate endophytic (DSE) association with a lulworthioid fungus (“*Lulwoana* sp.”), which is in conflict with several other studies from the NW Mediterranean Sea that point at the recently described pleosporalean fungus *Posidoniomyces atricolor*.
I collected *P. oceanica* roots at eight sites around Sicily and checked them for fungal colonization using light microscopy. At three sites, root fungal symbionts (=mycobionts) were isolated into pure cultures and identified using sequencing of the ITS rDNA gene.
*Posidoniomyces atricolor* represented the most frequent mycobiont (56 isolates), closely followed by lulworthioid fungi (51). The obtained mycobiont spectrum also comprised *Cladosporium* (2), *Alternaria* (1), *Corollospora* (1), *Fusarium* (1), *Penicillium* (1) and *Vishniacozyma* (1) isolates. The characteristic DSE root colonization similar to those occurring in terrestrial plants but not known from any other seagrass was found in all investigated *P. oceanica* individuals. The microscopic screening suggests that *P. atricolor* is indeed responsible for the observed DSE colonization.
This study extends the known range of *P. atricolor* and the DSE association characteristic for *P. oceanica* for southern Tyrrhenian Sea/Sicily. While lulworthioid fungi regularly occur in *P. oceanica* tissues, including terminal fine roots, their significance and functioning are unknown and beg further investigation. However, there are currently no proofs that they belong among dark septate endophytes of this seagrass.

**One-sentence summary:** This paper corrects an opinion that “*Lulwoana* sp.” (Lulworthiales) is a dark septate endophyte of the dominant Mediterranean seagrass *Posidonia oceanica*, because all available evidence suggests that the dark septate endophytic association typical for this seagrass is formed by its specific root mycobiont *Posidoniomyces atricolor* (Pleosporales).

## Introduction

Seagrasses are a taxonomically narrow but ecologically important group of vascular plants adapted to permanent life beneath the sea surface (Hemminga & Duarte 2000). Among the many adaptive traits, they take up mineral nutrients through the leaves, but under oligotrophic conditions, they rely on the roots and the nutrients present in the seabed substrate. While the vast majority of terrestrial vascular plants obtain nutrients through mycorrhizae, i.e., specialized root-fungus organs evolved to help the photoautotroph to scavenge minerals from recalcitrant substrates and the chemoheterotroph to access carbon and energy from photosynthates, seagrasses are regarded non-mycorrhizal (Nielsen *et al*. 1999). However, their roots do host fungal symbionts (=mycobionts), but their symbiotic modes (e.g., parasitic, pathogenic, endophytic) as well as ecophysiological roles, including a possible role in nutrient uptake, remain unknown.

*Posidonia oceanica* is the climax seagrass in the mostly oligotrophic Mediterranean Sea that is often characterized as a “marine desert” (Powley *et al*. 2017). Phosphorus and nitrogen are the most important limiting nutrients (Estrada 1996) and *P. oceanica* has to rely on their uptake through the roots (Lepoint *et al*. 2002). *Posidonia oceanica* typically colonizes coarse sandy to rocky bottoms, i.e., mostly mineral substrates, but with progressing time, it forms a compact substance composed of degrading leaves, rhizomes and roots mixed in various ratios with sediments and the seabed substrate, which is called matte. Matte can be several meters thick and hundreds to thousands of years old (Mateo *et al*. 1997) and stores significant amounts of organically-bound mineral nutrients (Romero *et al*. 1992). These are typically non-available to plant roots without the aid of heterotrophic organisms that decompose organic substrates and release mineral nutrients into the rhizosphere (mostly bacteria and non-symbiotic fungi) or transport them directly to the roots (mycorrhizal fungi). Characterization of the root mycobiota of *P. oceanica* is thus crucial for understanding of the seagrass’ mineral uptake and nutrient turnover in its rhizosphere.

One of the first attempts to document fungal colonization and cultivable mycobiont diversity of *P. oceanica* roots is the study by Torta *et al*. (2015). These authors visited two localities in NW Sicily, Italy and reported “inter- and intracellular septate mycelium, producing intracellular microsclerotia” that was “detected from the rhizodermis to the vascular cylinder” of the seagrass roots. However, the figures offered by Torta *et al*. (2015) contradict these claims; for example, the caption of Fig. 3a reads “transverse section with mycelium developing from the rhizodermis to the vascular cylinder” while Fig. 3a actually does not display any intraradical fungal mycelium. Additionally, they isolated a single mycobiont that was subsequently identified as “*Lulwoana* sp.”, based on a 99–100% ITS rDNA sequence similarity with the GenBank entry KC145432. The authors speculated that this mycobiont is a dark septate endophyte (DSE) and hypothesized that its presence “may help the host in several ways, particularly in capturing mineral nutrients [from rocks] through lytic activity”. However, there is no evidence available that the fungal structures depicted in Torta *et al*. (2015) actually belonged to this mycobiont, that this mycobiont is a DSE of *P. oceanica* nor that it could help the seagrass in any way. Indeed, the Lulworthiales mainly comprise saprobes inhabiting dead (drift-)wood, corals, herb stems and Foraminifera shells (Kohlmeyer *et al*. 2000) and despite that they are often recovered from marine algae and seagrasses, it is unknown whether such relationships are biotrophic and beneficial to these hosts. The lulworthioid genus *Lulwoana* currently contains only one described species (mycobank.org, accessed April 28, 2021), i.e., the lignicolous marine hyphomycete *L. uniseptata* (anamorph *Zalerion maritimum*) (Anastasiou 1963; Campbell *et al*. 2005). Nevertheless, based on the speculations by Torta *et al*. (2015), *Lulwoana* has already been cited as “fungal endophyte associated with seagrasses” (Supaphon *et al*. 2017), “fungal endophyte similar to dark septate endophytes” (Ettinger & Eisen 2019) and “known endophyte” (Kearns *et al*. 2019).

In the same year, Vohník *et al*. (2015) reported their observations from *P. oceanica* healthy terminal fine roots collected at several localities in the NW Mediterranean Sea. All their samples displayed a previously undocumented colonization pattern resembling terrestrial DSEs. Subsequently, identical colonization has been reported from dozens of other localities in the NW Mediterranean Sea (Vohník *et al*. 2017, 2019). In the following year, Vohník *et al*. (2016) reported the spectra of cultivable fungi isolated from the roots that had been used for the description of the DSE colonization. Surprisingly, they were dominated by a single unknown mycobiont with affinities to Aigialaceae (Pleosporales), with some other isolates belonging to the Lulworthiales. Similar scenario was repeated at many other localities in the NW Mediterranean Sea, also using high-throughput sequencing (Vohník *et al*. 2017, 2019) and the mycobiont was eventually described as *Posidoniomyces atricolor* (Vohník *et al*. 2019). There is convincing microscopic evidence that *P. atricolor* forms the DSE colonization detailed above (Vohník *et al*. 2019; Vohník 2021). At the same time, although ubiquitous in *P. oceanica* roots, this fungus has not been reported from any other host or environment. Taken together, these indications suggest a close specific biotrophic relationship of *P. atricolor* with the dominant Mediterranean seagrass.

There is a disagreement between the conclusions presented by Torta *et al*. (2015) and Vohník and colleagues (see above), mainly concerning the form and the distribution of the fungal colonization in *P. oceanica* roots. Therefore, I collected the seagrass’ roots at eight localities around Sicily and investigated them using light microscopy. At three localities, root mycobionts were isolated into pure cultures and identified using molecular tools. Due to the omnipresence of the DSE symbiosis/*P. atricolor* in the NW Mediterranean, I hypothesized that they would be present also at the Sicilian localities.

## Materials and Methods

### Investigated localities, sampling

I wanted to re-visit the sites investigated by Torta *et al*. (2015), but their precise location was not possible, because the provided coordinates are “301663E; 4228890N” for San Vito Lo Capo and “378638E; 4209008N” for San Nicola l’Arena. Therefore, I used the locality names combined with their approximate positions in the map presented in that paper and chose two accessible localities at San Vito Lo Capo (S. V. Lo Capo I/IT-78 and S. V. Lo Capo II/IT-79) and one accessible locality close to San Nicola l’Arena (as Trabia/IT-80) (Table 1, Suppl. Fig. 1). Root samples from these localities were used for screening of root fungal colonization by microscopy and mycobiont isolation and identification. In addition, five other localities around Sicily were visited (IT-73 to IT-77, Table 1, Suppl. Fig. 1) and the respective root samples were used only for microscopy.

**Table 1:** Localities on Sicily investigated in this study.

| locality name | locality ID | GPS coordinates | date of collection | collected by | microscopic screening | mycobiont isolation |
| --- | --- | --- | --- | --- | --- | --- |
| Milazzo | IT-73 | N38.23165 E15.24870 | 31.8.2015 | O. Borovec | yes | no |
| Capo Passero | IT-74 | N36.66941 E15.12696 | 1.9.2015 | M. Vohník | yes | no |
| Torre di Gaffe | IT-75 | N37.13954 E13.83171 | 2.9.2015 | M. Vohník | yes | no |
| Sciacca | IT-76 | N37.50342 E13.09337 | 2.9.2015 | O. Borovec | yes | no |
| Biscione | IT-77 | N37.69905 E12.47618 | 2.9.2015 | O. Borovec | yes | no |
| San Vito Lo Capo I<sup>1</sup> | IT-78 | N38.18363 E12.73246 | 3.9.2015 | O. Borovec | yes | yes |
| San Vito Lo Capo II<sup>1</sup> | IT-79 | N38.14307 E12.73683 | 3.9.2015 | M. Vohník | yes | yes |
| Trabia<sup>2</sup> | IT-80 | N37.99697 E13.65727 | 3.9.2015 | M. Vohník | yes | yes |

<sup>1</sup> these localities are identical to or in the immediate vicinity of those investigated by Torta et al. (2015) (see Materials and Methods)

Since endophytism is a biotrophic relationship and Torta *et al*. (2015) hypothesized a role of their mycobiont in “capturing mineral nutrients through lytic activity”, I focused on the part of the *P. oceanica* root system where one could expect the highest nutrient uptake, i.e., the healthy-looking turgescent terminal (=first order) roots (ca. 0.5–1 mm in diam.). They were collected at various depths and substrates (with the aim to include the shallowest and the deepest *P. oceanica* individuals/patches available as well as to reflect the respective seabed heterogeneity) using scuba diving in August/September 2015. At each locality, root samples were taken from five different *P. oceanica* individuals/patches at least 5 m apart, i.e., 40 individuals/patches were sampled in total. The samples for mycobiont isolation were processed within 24 hours from sampling whereas those for microscopy were immediately transferred to 50% ethanol and stored in a fridge (ca. 6°C) until processed.

### Root colonization

The microscopic screening and documentation of root fungal colonization followed Vohník *et al*. (2015). The intensity of fungal colonization was estimated based on the presence/absence of typical fungal extraradical and intraradical structures, i.e., superficial hyphae, superficial hyphal parenchymatous sheaths, intracellular hyphae in the rhizodermal cells and intracellular microsclerotia in the “entry cells” (Vohník *et al*. 2015) and the hypodermis. Ten hand-made longitudinal sections (ca. 3–5 mm long) from ten random roots were screened per each sampled individual, i.e., 400 sections were screened in total (Suppl. Table 1). No root staining was used. The root colonization was documented using an Olympus BX60 microscope equipped with DIC and an Olympus DP70 camera + QuickPHOTO MICRO 3.2 software (Promicra, Czechia). The Deep Focus mode (Promicra) was employed when beneficial. The photographs were modified for clarity as needed (adjustment of brightness and contrast) using Paint.NET ver. 4.0.13 (https://www.getpaint.net/).

### Mycobiont isolation and identification

Mycobionts were isolated from surface-sterilized root segments (ca. 3 mm in length) by incubation on potato dextrose agar at room temperature in the dark for 167 days. There were 30 root segments per each of the five individuals/patches per each of the three localities, i.e., 150 segments per locality and 450 segments in total.

PCR was performed using the ITS1F+ITS4 primer pair while the sequencing reaction using the ITS1, ITS1F and/or ITS4 primers. The ITS region of the rDNA was selected because Torta *et al*. (2015) targeted this gene, despite that phylogeny of the Lulworthiales is based rather on the SSU and LSU rDNA (Campbell *et al*. 2005; Azevedo *et al*. 2017). The isolates’ molecular identification followed Vohník (2020). Representative sequences of lulworthioid isolates were aligned with those from Torta *et al*. (2015), i.e., KF719964–9, and the resulting alignment was used for a delimitation of molecular operational taxonomic units (MOTUs) at 99% sequence similarity, following a NJ analysis with automatically estimated parameters in TOPALi (Biomathematiscs & Statistics Scotland). The alignment is available from the author upon reasonable request.

Colony and mycelium morphology of all representative isolates were documented using the same tools as described above for root colonization.

## Results

### Root colonization

The great majority of the terminal root sections displayed at least some fungal colonization and the average proportion of non-colonized sections across all localities was only 3% (min. 0%, max. 8%, Suppl. Table 1). At two localities (IT-79 and IT-80), all sections were colonized, in contrast to IT-78 where non-colonized sections reached the highest 8%. Intriguingly, IT-78 is the closest/identical locality to one of the localities investigated by Torta *et al*. (2015) (i.e., San Vito Lo Capo) and IT-79 lies in its close vicinity.

The average proportion of superficial hyphae (Fig. 1A, B) was high at all localities (min. 92%, max. 100%, avg. across all localities 97%) but the rest fluctuated considerably; the second highest were intracellular hyphae in the rhizodermis (Fig. 1C, D; 56%, 94%, 77.8%) followed by intracellular microsclerotia (Fig. 1E, F; 16%, 70%, 41%) and the least frequent superficial hyphal parenchymatous sheaths (Fig. 1G, H; 4%, 46%, 27%) (Suppl. Table 1).

No new colonization patterns were detected and no macroscopic sporocarps were visible on the surface of the screened roots. Five root segments displayed some infrequent root hairs with terminal swellings (Fig. 1K; max. one segment per locality). Several root hairs were cracked, revealing their spiral cell walls (Borovec & Vohník 2018; Kolátková & Vohník 2019).

**Figure 1:**
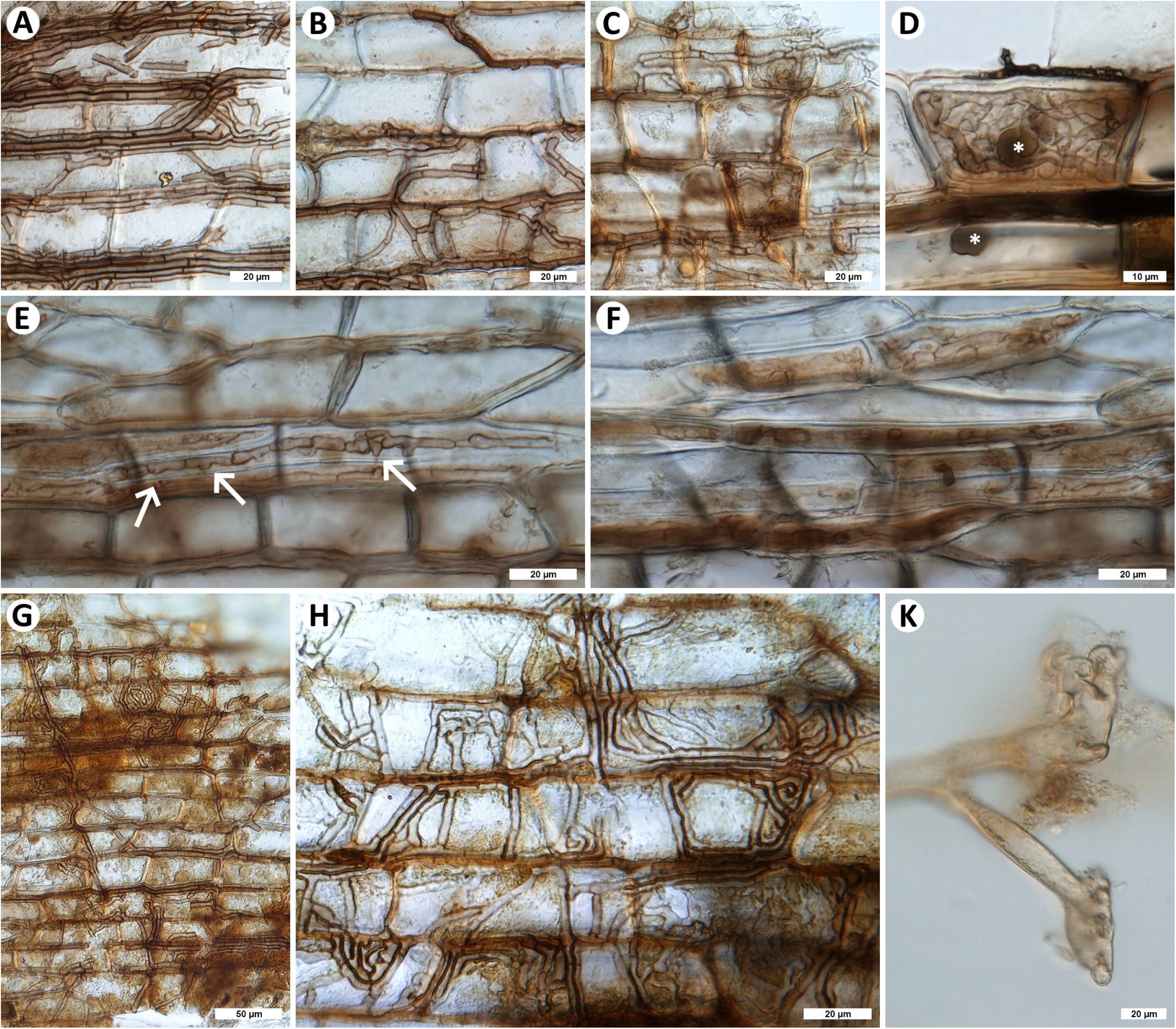
Dark septate endophytic colonization pattern in the roots of *Posidonia oceanica* from Sicily. **A, B:** superficial dark septate hyphae either growing mostly in parallel or following the sulci on the root surface/randomly. **C, D:** intracellular dark septate hyphae inside rhizodermal cells, note host nuclei in D (asterisks). **E, F:** early and late stages of microsclerotia development, note frequent connections between two parallel intracellular hyphae in E (arrows). **G, H:** superficial hyphal parenchymatous sheaths. **K:** terminal swellings at the end of a *P. oceanica* root hair. No staining was used, photos are from an upright microscope equipped with DIC, for more info see Materials and Methods. Bars A, B, C, E, F, H, K = 20 µm, D = 10 µm and G = 50 µm.

### Mycobiont isolation and identification

In total, 128 isolates were obtained, corresponding to a ca. 28% overall isolation success. However, there were apparent differences between the three investigated localities: IT-78 yielded 14 isolates (9.3%), IT-79 yielded 46 isolates (30.7%) and IT-80 yielded 68 isolates (45.3%) (Table 2).

**Table 2:** ***Posidonia oceanica* root mycobionts isolated in this study**

| locality ID<sup>1</sup> | isolation success<sup>2</sup> | total isolates | total identified isolates | total *Posidoniomyces atricolor* (representatives in GenBank)<sup>3</sup> | total *Lulworthiales* spp. (representatives in GenBank)<sup>3</sup> | total other identified mycobionts (representatives in GenBank)<sup>4</sup> |
| --- | --- | --- | --- | --- | --- | --- |
| IT-78 | 9.3% | 14 | 13 | 6/46%<br>(MW218888) | 6/46%<br>isolate SIC-121/MW218889<br>SIC-125/MW218891 | 1x *Penicillium* sp. (SIC-123/MW218890) |
| IT-79 | 30.7% | 46 | 37 | 7/19%<br>(MW218883) | 28/76%<br>SIC-093/MW218884<br>SIC-108/MW218886 | 1x *Cladosporium* sp. (SIC-107/MW218885) |
|  |  |  |  |  |  | 1x *Alternaria* sp. (SIC-110/MW218887) |
|  |  |  |  |  |  | (+7 unidentified bacteria/yeast-like) |
| IT-80 | 45.3% | 68 | 64 | 43/67%<br>(MW218882) | 17/27%<br>SIC-005/MW218878<br>SIC-021/MW218879 | 1x *Corollospora maritima* (SIC-023/MW218880) |
|  |  |  |  |  |  | 1x *Vishniacozyma victoriae* (SIC-025/MW218881) |
|  |  |  |  |  |  | 1x *Fusarium* sp. (not submitted) |
|  |  |  |  |  |  | 1x *Cladosporium* sp. (not submitted) |
|  |  |  |  |  |  | (+3 unidentified bacteria/yeast-like) |
| Average/total | 28.4% | 128 | 114 | 56 (49.1%) | 51 (44.7%) | 7 (+10 unidentified bacteria/yeast-like) |

<sup>1</sup> locality IDs identical to Table 1
<sup>2</sup> out of 150 surface sterilized *Posidonia oceanica* fine root segments per each locality (i.e., 30 segments per each sampled individual/patch, see Materials and Methods).
<sup>3</sup> the values after slashes represent % frequency among all identified isolates
<sup>4</sup> the unidentified bacteria/yeast-like isolates were not included in the total number of isolates nor in the subsequent calculations

Ten isolates represented unidentified yeasts. Out of the remaining 118 mycelial forms, 114 were identified (Table 2): 56 isolates (49.1%) represented *Posidoniomyces atricolor* and 51 isolates (44.7%) belonged to lulworthioid fungi. There were apparent differences in the recovery of these two mycobiont guilds at two localities: while IT-79 yielded 7 isolates of *P. atricolor* (19%) and 28 lulworthioid isolates (76%), IT-80 yielded 43 isolates of *P. atricolor* (67%) and 17 lulworthioid isolates (27%). Other isolated mycobionts comprised (alphabetically) *Alternaria* sp., *Cladosporium* spp., *Corollospora maritima, Fusarium* sp., *Penicillium* sp. and *Vishniacozyma victoriae* (syn. *Cryptococcus victoriae*). Sequences of representative isolates are deposited in GenBank at NCBI (https://www.ncbi.nlm.nih.gov/genbank) under the accession numbers MW218878–91 (Table 2).

At the 99% sequence similarity threshold, the lulworthioid fungi obtained in this study were divided into two MOTUs: the majority grouped with the “*Lulwoana* sp.” sequences KF719964, KF719966 and KF719967 from Torta *et al*. (2015) while two isolates (SIC-121/MW218889 and SIC-125/MW218891, Table 1) formed a separate MOTU. In addition, KF719965, KF719968 and KF719969 from Torta *et al*. (2015) formed another separate MOTU (data not shown). However, whether these MOTUs represent different lulworthioid species cannot be decided based only on ITS rDNA sequences.

All the isolated mycelia were septate and all the representative isolates comprised melanized (light to dark brown) hyphae, especially in later ontogenetic stages (=older colonies) (Fig. 2). On the other hand, only *P. atricolor* colonies originated directly from the intracellular microsclerotia formed inside the *P. oceanica* root segments (Fig. 2E, F).

**Figure 2:**
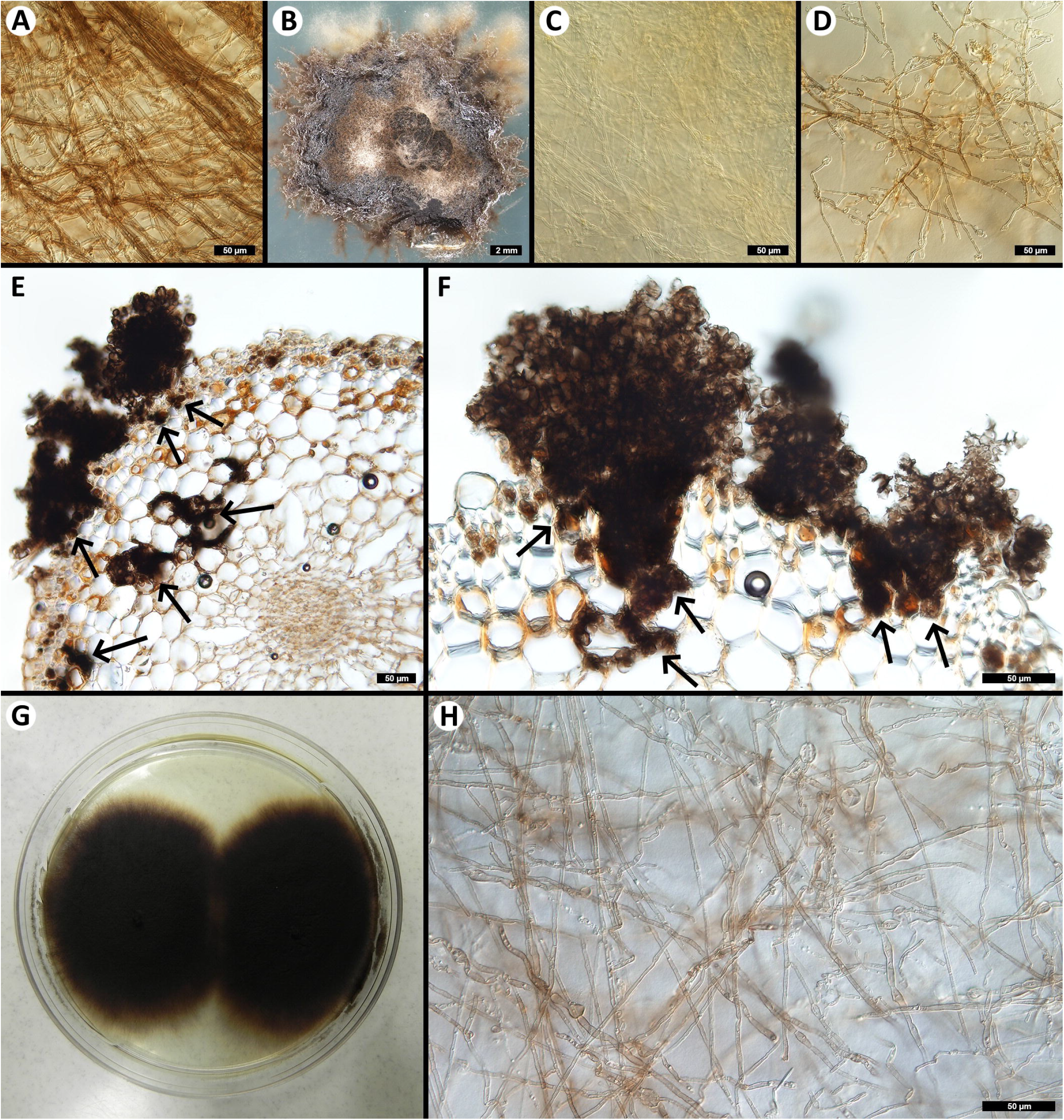
Colonial and hyphal morphology and anatomy of representative fungal isolates obtained in this study. **A:** *Cladosporium* sp. (isolate SIC-026) hyphae. **B:** Lulworthiales sp. (SIC-100) colony morphology and color. **C:** Lulworthiales sp. (SIC-100) young hyphae. **D:** Lulworthiales sp. (SIC-100) older hyphae. **E, F:** Transverse sections through a *P. oceanica* root segment with *Posidoniomyces atricolor* colonies on its surface, developing *in vitro* from intracellular darkly pigmented microsclerotia (arrows). Note that the intraradical fungal colonization is restricted to the upper layers of the cortex (incl. the hypodermis) and the rhizodermis, while never reaching the stele. **G:** *Vishniacozyma victoriae* (syn. *Cryptococcus victoriae*) (SIC-025) old colony morphology and color. **H:** *Vishniacozyma victoriae* (SIC-025) young mycelium. Bars A, C, D, E, F, H = 50 µm and B = 2 mm, the Petri dish in G has 8.5 cm in diam.

## Discussion

In contrast to Torta *et al*. (2015) and in agreement with several previous studies from the NW Mediterranean Sea (see above), I found the typical DSE colonization pattern in *Posidonia oceanica* roots at all localities investigated in this study. Likewise, *Posidoniomyces atricolor* had not been isolated by Torta *et al*. (2015) but represented up to 67% (average 49%) of the isolates identified in this study. The most likely reasons for this difference are the second-order roots investigated by Torta *et al*. (2015) vs. the first-order roots investigated here and especially the too short cultivation time (four weeks) employed by Torta *et al*. (2015). Indeed, many obligate marine fungi are notoriously slow growing (up to two years of incubation may be needed for some species, see Kohlmeyer *et al*. (2004)) and this is also true for *P. atricolor* whose colonies start to emerge from *P. oceanica* root segments only after ca. 1.5–2 months of incubation (Vohník *et al*. 2019).

Comparing the diversity of fungal isolates obtained in Torta *et al*. (2015) and here is at least to some extent complicated by the identification approach presented in the former, i.e., targeting the ITS rDNA (cf. Campbell *et al*. 2005; Azevedo *et al*. 2017) and relying on the first BLAST hit in GenBank at NCBI (cf. Nilsson *et al*. 2012). Nevertheless, as far as the ITS rDNA sequences can tell, probably only one lulworthioid species had been isolated by Torta *et al*. (2015) and the same species was detected at all localities investigated here. In addition, probable members of seven other genera were isolated here, suggesting that the seagrass’ root mycobiota at the Sicilian localities is not only richer than as reported by Torta *et al*. (2015) but the richest ever reported in studies employing mycobiont cultivation from surface sterilized *P. oceanica* root segments (Vohník *et al*. 2016, 2017; Vohník 2021).

Similarly to some other studies from the NW Mediterranean (see above), *P. atricolor* colonies were found to develop from intracellular darkly pigmented microsclerotia that occurred in the upper layers of the root cortex, suggesting that this mycobiont is indeed responsible for the observed DSE colonization pattern. Torta *et al*. (2015) reasoned that their “*Lulwoana* sp.” is a DSE, because it formed slow growing colonies “dark in color, with septate mycelium”. However, all representative isolates obtained in this study, including ubiquitous opportunistic saprobes also known from terrestrial ecosystems, produced dark septate hyphae in pure culture. This underscores the fact that DSE cannot be recognized based on the appearance of their colonies/mycelium formed outside of the host roots/under in vitro conditions. Instead, they are defined by morphology and color of the structures they produce in the host, i.e., relatively thick melanized septate hyphae colonizing the root surface and forming intracellular resting and storage organs known as microsclerotia (Stoyke & Currah 1991; Lukešová *et al*. 2015).

To conclude, this study extends the known range of *P. atricolor* for southern Tyrrhenian Sea/Sicily and provides additional support for the hypothesis that this mycobiont is responsible for the typical DSE colonization pattern observed in the roots of *P. oceanica*. It also confirms that *P. oceanica* root mycobiota regularly comprises related lulworthioid fungi, but their significance and functioning remain unknown. Unlike *P. atricolor*, most of them are relatively easy to culture so they could be used in resynthesis experiments to reveal their actual colonization pattern, possibly with a model non-seagrass hosts, since many endophytes do not seem to be host specific. However, the experimental results available to date indicate negative effects of lulworthioid fungi on their host plants (Hughes *et al*. 2020).

## Acknowledgments

This work was supported by the Institute of Botany, Czech Academy of Sciences (RVO 67985939). I wish to thank my former student Ondřej Borovec for the help with the sampling on Sicily as well as with the *P. oceanica* root mycobiont isolation and identification, Jiří Machač for assembling Figs 1 and 2 and Paula A. Buil Maldonado for the suggestions on an earlier version of this paper.

## FIGURE LEGENDS

**Supplementary Figure 1: Localities on Sicily investigated in this study**

*Posidonia oceanica* root samples were collected in August 2015 at eight localities around Sicily, Italy (IT-73 to IT-80, red circles; locality codes explained in Table 1) and all were checked for fungal colonization using light microscopy. In addition, cultivable fungi associated with the roots were isolated from samples from three localities (IT-78 to IT-80, black dots in red circles). The obtained results were compared with (Torta *et al*. 2015) whose localities are marked with asterisks. For more details see Materials and Methods. Bar = 20 km, the map was modified from OpenStreetMap (OpenStreetMap Foundation, for copyright info see https://www.openstreetmap.org/copyright).

