## Supplementary Table 1 for "No evidence that lulworthioid fungi are dark septate endophytes in the roots of the dominant Mediterranean seagrass *Posidonia oceanica*"

**Supplementary Table 1: Root fungal colonization in *Posidonia oceanica* individuals/patches investigated in this study**

| **locality ID^1^ (n=50)** | **average presence of^2^** | | | | **roots without any colonization** |
| --- | --- | --- | --- | --- | --- |
|  | **superficial hyphae** | **superficial hyphal parenchymatous sheaths** | **intracellular hyphae in the rhizodermal cells** | **intracellular microsclerotia** |  |
| IT-73 | 96% (90–100) | 30% (10–60) | 60% (30–70) | 56% (30–70) | 2% (0–10) |
| IT-74 | 98% (90–100) | 24% (10–40) | 76% (50–90) | 40% (10–80) | 2% (0–10) |
| IT-75 | 96% (80–100) | 4% (0–10) | 70% (10–100) | 22% (0–40) | 4% (0–20) |
| IT-76 | 96% (80–100) | 22% (0–70) | 56% (20–100) | 16% (0–40) | 4% (0-20) |
| IT-77 | 98% (90–100) | 46% (30–70) | 92% (80–100) | 30% (20–50) | 2% (0–10) |
| IT-78 | 92% (90–100) | 20% (0–60) | 80% (60–100) | 24% (10–30) | 8% (0–10) |
| IT-79 | 100% | 34% (10–60) | 94% (90–100) | 70% (60–80) | 0% |
| IT-80 | 100% | 32% (20–40) | 94% (90–100) | 68% (40–100) | 0% |
| *All localities* | *97%* | *27%* | *77.8%* | *41%* | *3%* |

^1^ locality IDs identical to Table 1

^2^ calculated based on screening of 10 longitudinal sections, each from one of the 10 roots per each of the 5 sampled *P. oceanica* individuals/patches per each locality (for details see Materials and Methods). The values are means followed by range (minimum–maximum) except the last row where only means are given.
