## Supplementary figures and images for "No evidence that lulworthioid fungi are dark septate endophytes in the roots of the dominant Mediterranean seagrass *Posidonia oceanica*"

### Supplementary Figure 1

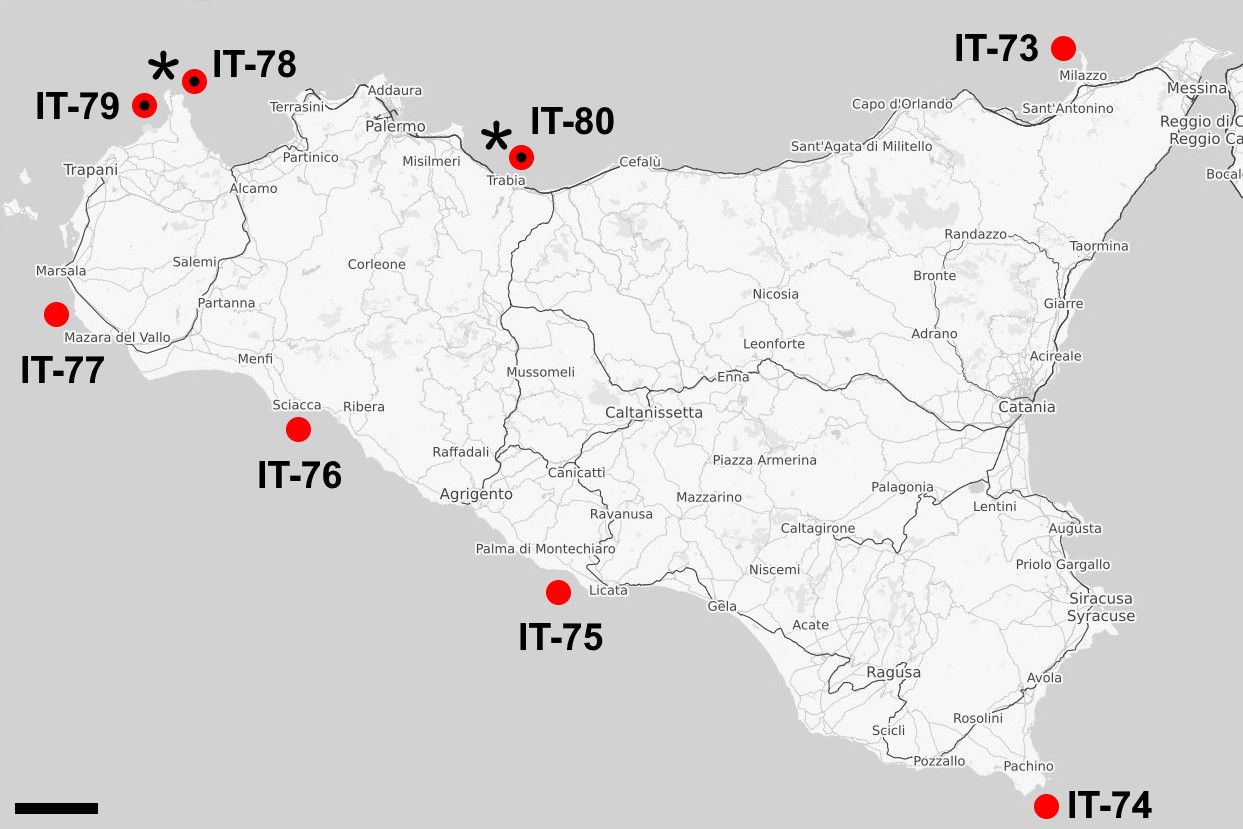
